# Effect of temperature on life-history traits and mating calls of a field cricket, *Acanthogryllus asiaticus*

**DOI:** 10.1101/2020.06.06.137869

**Authors:** Richa Singh, P Prathibha, Manjari Jain

## Abstract

Ectotherms are sensitive to the changes in ambient temperature with respect to their physiology and development. To compensate for the effects of variation in temperature, ectotherms exhibit physiological plasticity which can be for short or long term. An extensive body of literature exists towards understanding these effects and the solutions ectotherms have evolved. However, to what extent rearing temperature during early life stages impacts the behaviour expressed in adulthood is less clearly understood. In the present study, we aimed to examine the effect of developmental temperature on life-history traits and mating call features in a tropical field cricket, *Acanthogryllus asiaticus*. We raised *A. asiaticus* at two different developmental conditions: 25°C and 30°C. We found developmental time and adult lifespan of individuals reared at 30°C to be shorter than those at 25°C. Increased developmental temperature influenced various body size parameters differentially. Males raised at 30°C were found to be larger and heavier than those raised at 25°C, making *A. asiaticus* an exception to the temperature-size rule. We found a significant effect of the change in immediate ambient temperature on different call features of both field-caught and lab-bred individuals. In addition, developmental temperature also affected mating call features as individuals raised at higher temperature produced faster calls with a higher peak frequency compared to those raised at lower temperature. However, the interaction of both developmental and immediate temperature on mating calls showed differential effects. Our study highlights the importance of understanding how environmental temperature shapes life-history and sexual communication in crickets.

## 1. Introduction

Temperature is a crucial environmental factor which affects all organisms in myriad ways, particularly ectotherms, given that they do not maintain constant body temperature and most of their physiological functions are regulated by ambient temperature (Bartholomew and Tucker, 1963). Individuals in their natural environment often face varying temperature regimes through their developmental stages to adulthood. Such changes in temperature affect their behaviour, growth (change in mass), development (change in morphology) and physiology with significant impacts on life-history and fitness (Nylin and Gotthard, 1998; Abram et al., 2017). However, in response to the changes in environmental temperature, ectotherms can exhibit physiological adjustment (plasticity/acclimation) to maintain their performance and fitness in an altered environment (Wilson and Franklin, 2002; Seebacher et al., 2015). Acclimation is achieved by shifting the temperature for optimum performance (thermal sensitivity) of various physiological and biochemical reactions (Guderley and Pierre, 2002). For example, in common carp (*Cyprinus carpio*), locomotor performance is maintained in different environmental temperatures by shifting the thermal sensitivity of different enzymes (Johnston and Temple, 2002). Ectotherms can show developmental (or phenotypic) plasticity, which is the ability of a genotype to produce different phenotypes in response to different environmental conditions (West-Eberhard, 2003). Thus, the temperature during developmental stages can influence behaviour in adulthood. For example, in the European honey bees (*Apis mellifera*), a colony raised at a relatively low temperature had a delayed onset of foraging and fewer dancers due to the effect of developmental temperature on hormone metabolism (Becher et al., 2009). In the field cricket, *Gryllus bimaculatus*, individuals raised at high temperatures were more explorative and had a lower coefficient of variation of the behaviour within individuals for all the temperature treatments (Niemelä et al., 2019).

Environmental temperature can act as a critical determinant of various life-history traits such as longevity (Bauerfeind et al., 2009), developmental time (Ciota et al., 2014) and adult size (Atkinson, 1994) in insects. In insects, developmental time (the time taken to reach adulthood) generally decreases with increasing temperature as observed in the three species of *Culex* mosquitoes (Ciota et al., 2014). The model by Gillooly et al. (2001), explains the mechanism of this trend as it predicts that an increase in temperature will increase the metabolic rate, which reduces the developmental time. In most ectotherms, individuals reared at low temperatures take longer to develop but have larger bodies at equivalent developmental stages than those reared at high temperatures, referred to as the temperature-size rule (Atkinson, 1994). This rule is a particular case of Bergmann’s rule, which signifies that the relationship between environmental temperature and body size is the product of phenotypic plasticity (Atkinson, 1996). The biophysical model proposed by Van der Have and De Jong (1996) suggests that the higher activation energy or temperature for differentiation compared to growth is the mechanistic explanation for the temperature-size rule. However, the general reason for this rule remains elusive and scientists have expressed doubt on the applicability of this rule to ectotherms (Walters and Hassall, 2006).

Environmental temperature can also affect various behaviours of ectotherms, such as defensive behaviour (Passek and Gillingham, 1997), foraging behaviour (Le lann et al., 2011) and mating behaviour (Brandt et al., 2018). In the context of mating behaviour, temperature influences different parameters of sexual signals such as the amplitude and frequency of electric discharge of knifefish, *Apteronotus leptorhynchus* (Dunlap et al., 2000). The effect of immediate ambient temperature on the acoustic properties of the mating signal is particularly well studied in acoustically active insects (Martin et al., 2000; Hedrick et al., 2002; Greenfield and Medlock, 2007) and anurans (Gerhardt, 1978; Zweifel, 1968). However, the effect of developmental temperature on mating signals is not well understood. In crickets, sound is produced by the stridulation of modified forewing. It is expected that their calls will also vary with temperature, as the neuromuscular system which is involved in sound production gets affected by the variation in temperature (Martin et al., 2000; Walker and Cade, 2003). Dolbear (1897) reported the utility of cricket chirp rates as a thermometer since it increases linearly with temperature. While a plethora of studies has examined the effect of immediate ambient temperature on the intersexual acoustic signals of crickets, few studies have examined the effect of developmental temperature on them. To our knowledge, only three studies so far have examined the effect of developmental temperature on mating calls of crickets, *Allonemobius fasciatus* (Olvido and Mousseau, 1995), *Laupala cerasina*, (Grace and Shaw, 2004) and *G. rubens* (Beckers et al., 2019). Of these only the first examined the interactive effect of immediate ambient and developmental temperature on intersexual call features.

*Acanthogryllus asiaticus* (family: Gryllidae) is a tropical field cricket, native to the Indian subcontinent (Gorochov, 1990). *A. asiaticus* is bivoltine and mostly active during the summer with a minor peak during the post-monsoon season in India (Singh and Jain, 2020). Males produce a stereotypic long-distance mating call (LDMC) with a relatively low peak frequency (4687.610 ± 482.08 Hz) to attract females (Singh and Jain, 2020). Since this is a bivoltine species, the two populations in a year face different temperature regimes during their development, making it an ideal system to study the effect of developmental temperature on life-history and mating calls. Hence, in this study, we examined the effect of developmental temperature on various life-history traits of *A. asiaticus* and the independent and interactive effect of immediate ambient temperature and developmental temperature on the properties of their long-distance mating calls.

## 2. Materials and methods

The study was carried out between April 2015 and December 2019. Adult males and females of *A. asiaticus* were collected from Indian Institute of Science Education and Research (IISER) campus in Mohali (30°39◻N, 76°43◻E) to set-up the laboratory culture of the species. In addition, field-caught adult males were also used for assessing the effect of immediate ambient temperature on mating call features.

### 2.1. Animal Husbandry

Ten mating pairs of lab-reared *A. asiaticus* were set for mating at 25°C. Eggs from each set were segregated and were equally divided into two sets, with one set kept in a room maintained at 25°C and the other in a climatic chamber (Memmert GmbH+Co.KG, Germany) maintained at 30°C. In both cases, relative humidity ranged from 40-70% and a daily 12L:12D light cycle was maintained. Individuals were reared and separated into individual boxes on reaching adulthood (for details, see Appendix S1).

### 2.2. Effect of temperature on life-history traits

With respect to life-history traits, developmental time (total number of days taken by nymphs to reach adulthood), adult lifespan (total number of days an individual survived after the final moult), and nymph appearance duration were measured for individuals from both the sets. Body morphometry of adults (for both males and females) was carried out for the following parameters: body length, pronotum length, pronotum width, wing size and ovipositor length for females. All the morphometric measurements were done with Leica stereo zoom microscope (M 205C, Leica Microsystems GmbH, Wetzlar, Germany) with an attached digital camera (Leica MC120HD, Leica Microsystems GmbH, Wetzlar, Germany) using LAS software V4.8 (Leica Microsystems, Switzerland). Bodyweight was measured using a weighing balance (Sartorius analytical balance: BSA224S-W, Sartorius AG, Goettingen, Germany).

### 2.3. Effect of temperature on mating call features

Animals were kept in individual plastic containers (diameter - 12 cm and height - 6 cm) covered with cloth mesh and were placed in a dark, quiet room (ambient noise at 15 dB at 5 kHz) maintained at the relevant recording temperatures for at least 5 hours prior to the recording to ensure acclimatisation. Post-acclimatisation audio recordings of calling males were made as 16-bit WAV files at a sampling rate of 44.1 kHz using Tascam, linear PCM recorder (DR-07 Mk II, TEAC Professional, USA). All recordings were digitised and analysed in Raven Pro1.5 (Cornell Laboratory of Ornithology, Ithaca, NY) to quantify the following temporal and spectral features: chirp duration, chirp period, syllable duration, syllable period, number of syllables per chirp and peak frequency.

Twenty adult males collected from the field were housed in individual boxes in the lab at 24°C, 40 - 70% humidity, 12L:12D light cycle. *Ad libitum* food and water were provided. After a week of acclimatisation, individuals were recorded one at a time, at five different ambient temperatures: 22°C (N = 7), 24°C (N = 9), 26°C (N = 5), 28°C (N = 7) and 30°C (N = 10). To examine the effect of immediate ambient temperature and developmental temperature on lab-bred individuals, animals were raised at two different temperatures 25°C (N = 12) and 30°C (N = 16) and recorded at either 25°C or 30°C on different nights.

### 2.4. Statistical analyses

Statistical tests were performed using Statistica 64 (Dell Inc.2015, Version 12). Shapiro-Wilk test was used to check for normality. Comparisons for examining the effect of developmental temperature on nymph appearance duration, developmental time, adult lifespan and body morphometric traits on individuals bred at 25°C and 30°C were done using t-tests. Effect of immediate ambient temperature on calling behaviour of field-caught individuals was tested using a Kruskal-Wallis test followed by pair-wise comparisons using the Mann-Whitney U test with Bonferroni corrections. Independent and interactive effect of immediate temperature and developmental temperature on various call features of lab-bred individuals was compared using a factorial multivariate analysis of variance (MANOVA). Here, immediate temperature and developmental temperature were taken as independent variables; 25°C versus 30°C as a categorical factor and all 7 call features were taken as dependent variables. In addition, a two-way analysis of variance tests was carried out individually for all call features to test for the independent and interactive effect of immediate ambient and developmental temperatures on call features. We also carried out pair-wise comparisons using t-tests to compare the effect of two temperatures (25°C and 30°C) on each call feature for both immediate and developmental temperature.

## 3. Results

### 3.1. Effect of temperature on life-history traits

Developmental temperatures were found to have a significant effect on different life-history traits (Fig. 1, Table S1). Nymphs hatched faster at 30°C than at 25°C (t-test, t = 20.57, df = 271, P < 0.01). Around 23% of nymph hatched at 25°C and 30°C, of which only 20% and 18% survived at 25°C and 30°C, respectively. Individuals raised at 30°C showed shorter developmental time (96 days) and reached adulthood faster than those at 25°C (171 days) (t-test, t = 10.29, df = 52, P < 0.01). A significant difference was found for adult lifespan between 25°C and 30°C, as individuals raised at 25°C lived longer than those at 30°C (t-test, t=2.53, df = 52, P = 0.014). Adult males raised at 30°C were found to have higher pronotum length, pronotum width and body weight compared to those raised at 25°C (t-test, P < 0.05; Fig. 2, Table S2) whereas no such differences were observed in females. However, females raised at 25°C were found to have a larger ovipositor length, but a smaller body length and wing size compared to those raised at 30°C (t-test, P < 0.05; Fig. 2, Table S2).

**Fig. 1.**
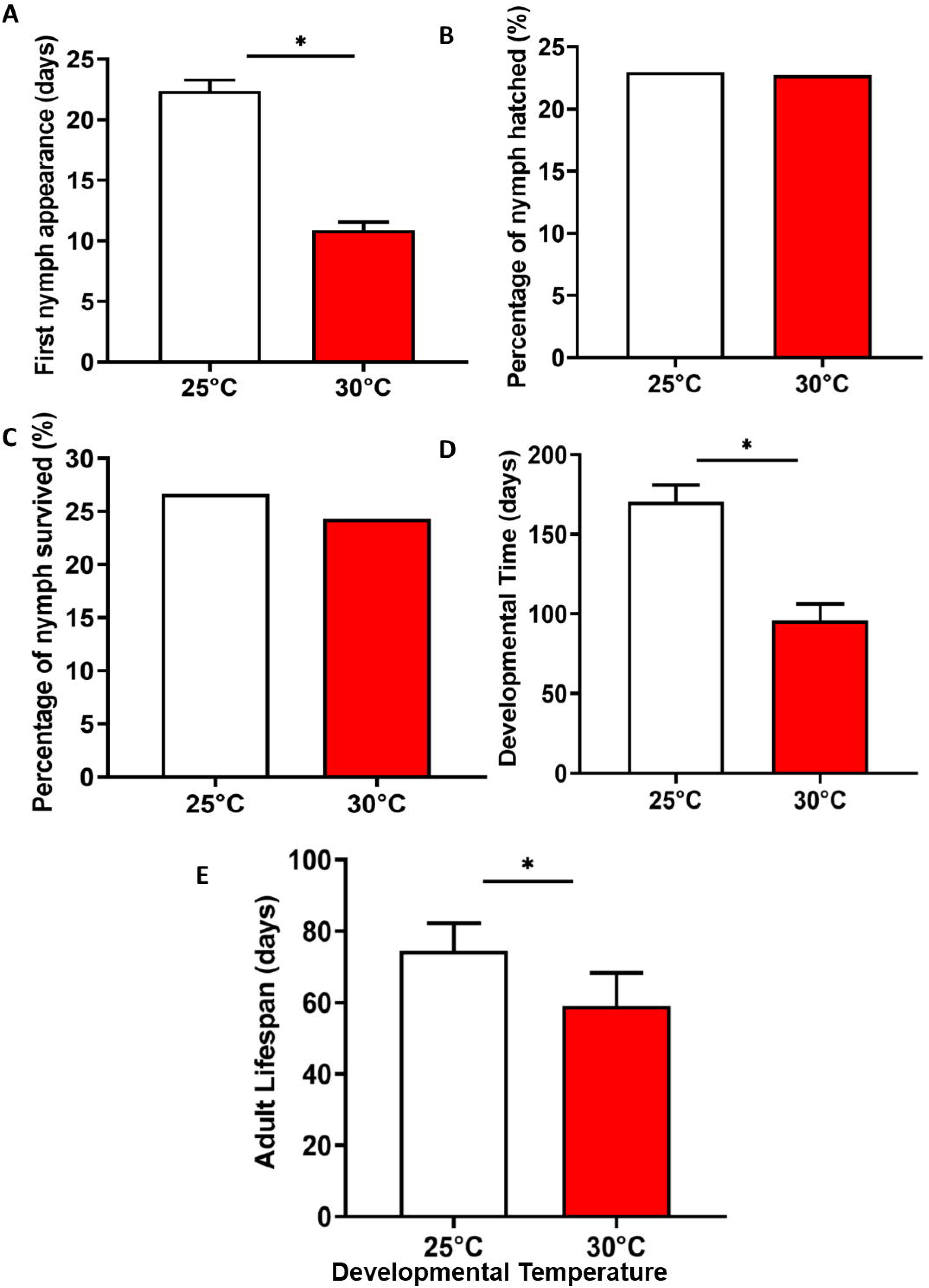
Comparison of effects of developmental temperatures: 25°C and 30°C on life-history traits of *Acanthogryllus asiaticus*. * indicates significant differences and ns indicates no significant difference. Mean ± 95% CI.

**Fig. 2.**
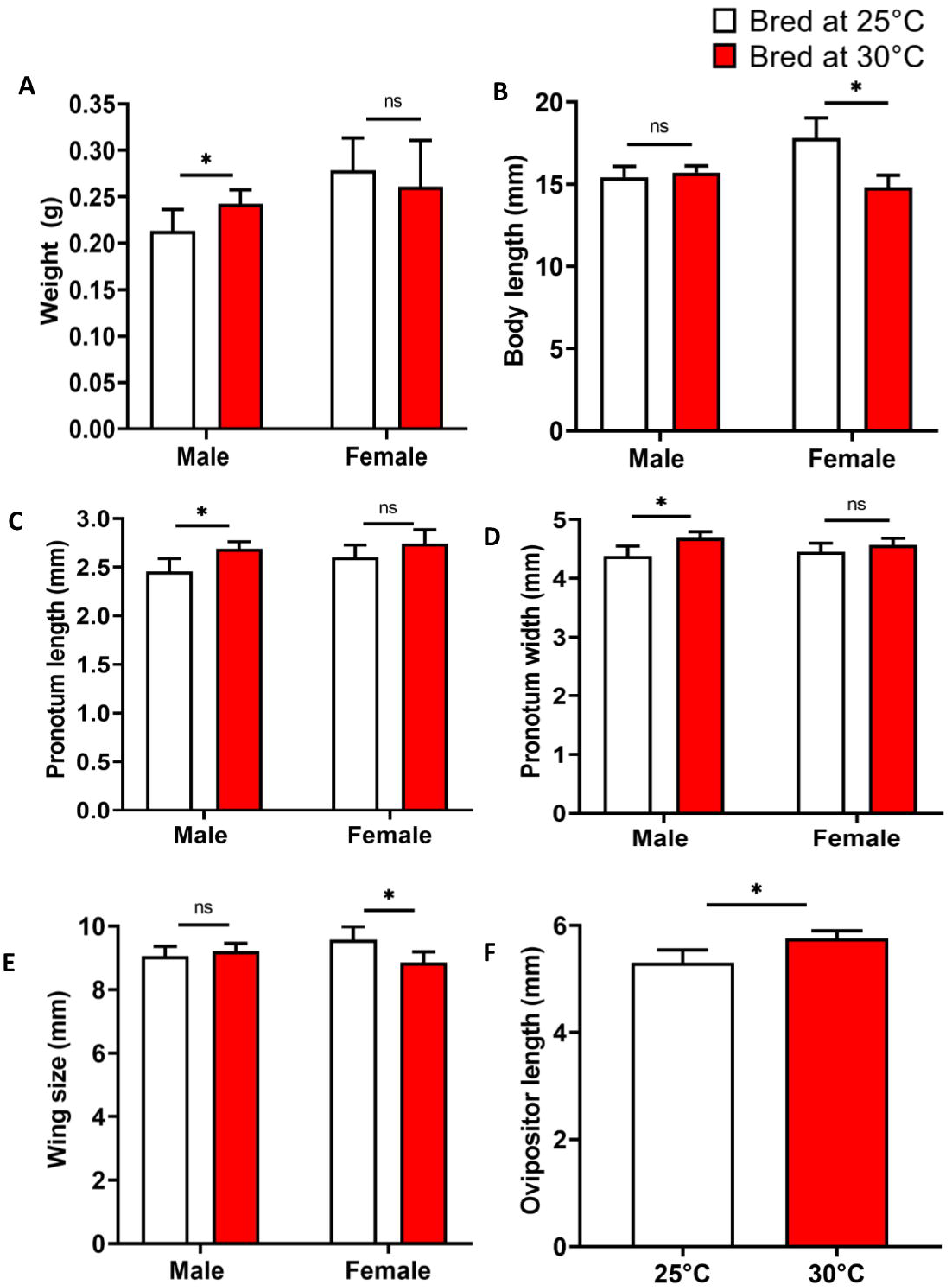
Comparison of effects of developmental temperatures: 25°C and 30°C on various body size parameters in males and females of *Acanthogryllus asiaticus*. * indicates significant differences and ns indicates no significant difference. Mean ± 95% CI.

### 3.2. Effect of temperature on mating call features

Immediate ambient temperature significantly impacted both temporal and spectral features of the mating call of field-caught animals (Kruskal-Wallis test, Fig. 3, Table S3). Chirp rate and peak frequency were found to increase while other call features such as chirp duration, syllable duration, number of syllables per chirp, syllable period were found to decrease with the rise in temperature (Mann-Whitney U test, Fig. 3, Table S3). Immediate temperature also influenced all call features except the number of syllables per chirp for lab-bred individuals (t-test, Fig.4, Table S4 & S5). Chirp period and chirp duration were found to decrease with temperature while chirp rate and peak frequency were found to increase with temperature.

**Fig. 3.**
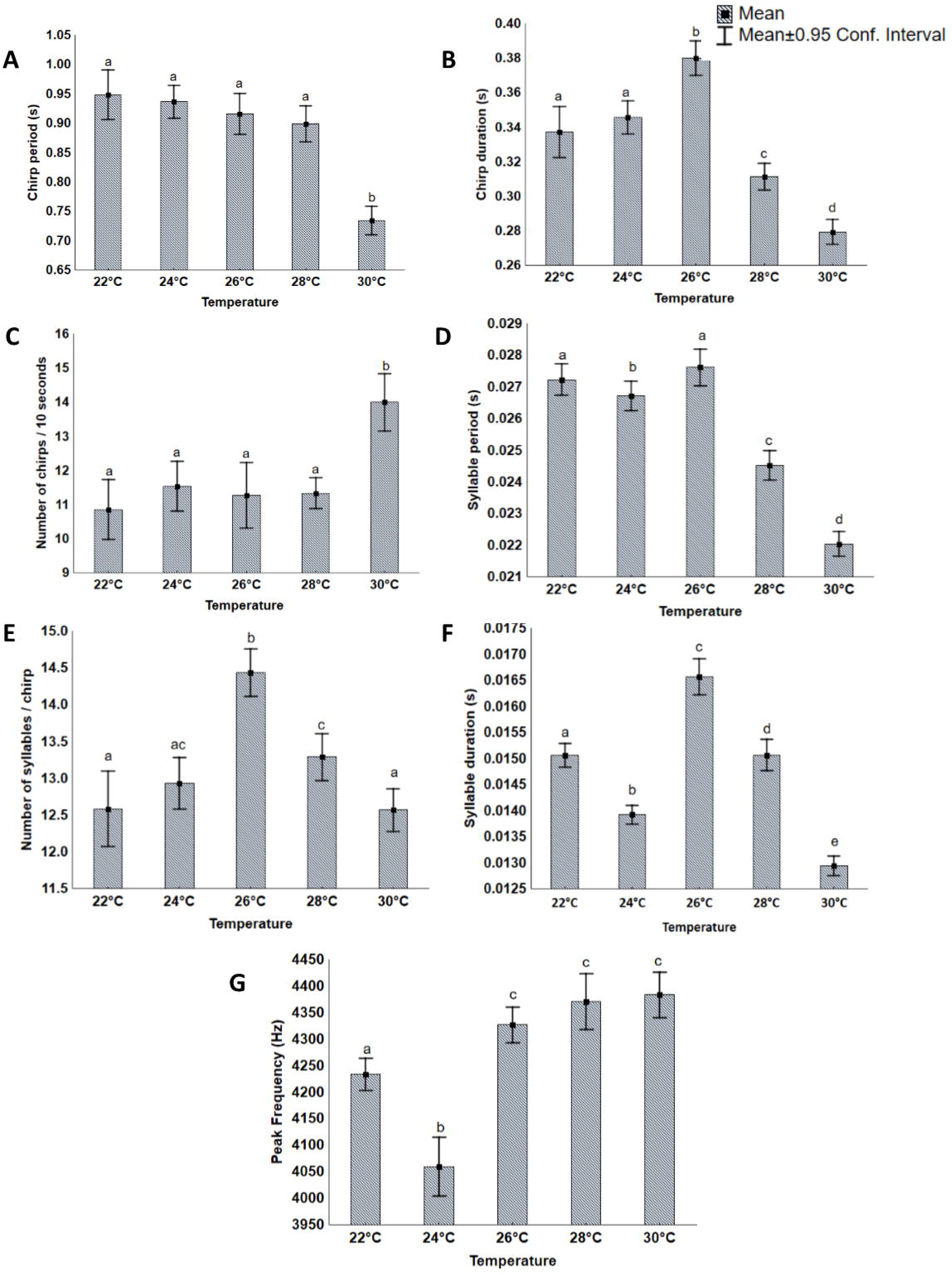
Effect of immediate temperature on different call features. Different letters indicate significant difference P < 0.05. Mean ± 95% CI.

**Fig. 4.**
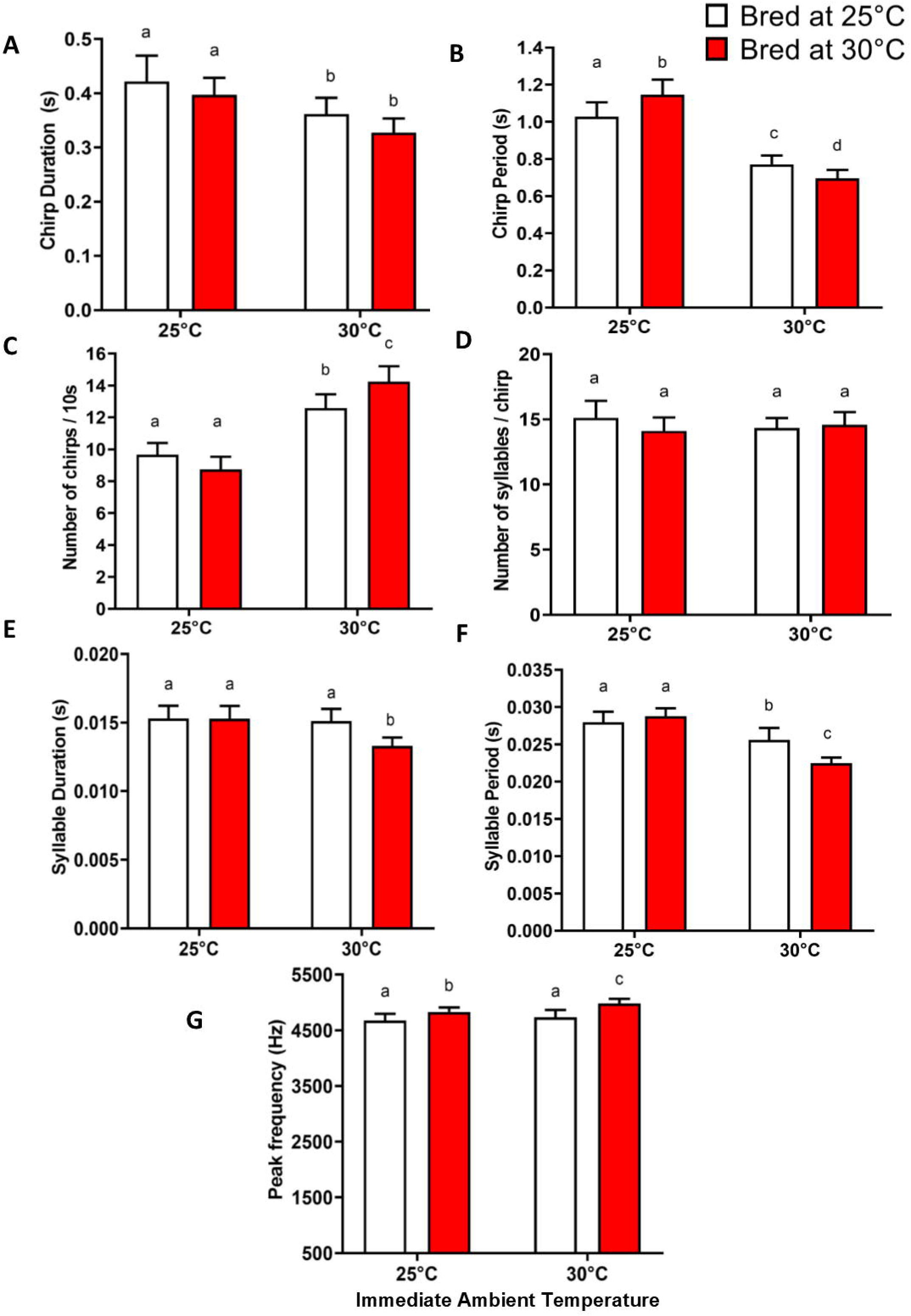
Effect of developmental temperature on different call features. Developmental temperatures: 25°C and 30°C; Recorded at ambient temperatures: 25°C and 30°C. Different letters indicate significant difference.

Factorial MANOVA showed independent effects of both developmental temperatures (Wilk’s λ = 0.562, F = 5.12, P < 0.01), immediate temperature (Wilk’s λ = 0.237, F = 21.11, P < 0.01) and a significant interactive effect of the two (Wilk’s λ = 0.646, F = 3.6, P < 0.01) on mating call features (Fig. 4, Table 1). Results of two-way analysis of variance conducted on individual call features demonstrated a significant independent effect of immediate temperature on all the call parameters except for the number of syllables per chirp. However, the independent effect of developmental temperature was found only on syllable duration, syllable period and peak frequency. Further, a significant interactive effect of immediate ambient temperature and the developmental temperature was found on all call features except peak frequency, chirp duration and number of syllables per chirp (Fig. 4, Table 1). Pair-wise comparisons revealed that chirp period, chirp rate, syllable duration, syllable period and peak frequency were significantly different between individuals bred at 25°C and 30°C, and recorded at 30°C (t-test, P < 0.05, Fig. 4, Table S6). However, a significant difference was only observed in the chirp period and peak frequency when individuals bred at 25°C and 30°C were recorded at 25°C, (t-test, P < 0.05, Fig. 4, Table S7).

**Table 1.** **A.** Multivariate analysis of variance (MANOVA) to examine the effect of immediate temperature and developmental temperature on the mating call of *Acanthogryllus asiaticus*. **B.** Effect of immediate temperature (25°C and 30°C) and developmental temperature (25°C and 30°C) on individual call features. 12 individuals bred at 25°C and 16 individuals bred at 30°C were recorded at 25°C and 30°C.

| **A** | **Wilks Value** | **F** | **df** | **p** |  |
| --- | --- | --- | --- | --- | --- |
| Intercept | 0.001 | 16983.01 | 7 | <0.01 |  |
| Developmental temperature | 0.562 | 5.12 | 7 | <0.01 |  |
| Immediate temperature | 0.237 | 21.11 | 7 | <0.01 |  |
| Developmental temperature*Immediate temperature | 0.646 | 3.6 | 7 | <0.01 |  |
| **B** | **Degree of Freedom** | **SS** | **MS** | **F** | **P** |
| **Chirp duration (s)** |  |  |  |  |  |
| Intercept | 1 | 7.787 | 7.787 | 2305.673 | < 0.01 |
| Developmental temperature | 1 | 0.012 | 0.012 | 3.528 | 0.066 |
| Immediate temperature | 1 | 0.058 | 0.058 | 17.054 | < 0.01 |
| Developmental temperature*Immediate temperature | 1 | 0.000 | 0.000 | 0.095 | 0.759 |
| Error | 52 | 0.176 | 0.003 |  |  |
| **Chirp period (s)** |  |  |  |  |  |
| Intercept | 1 | 45.449 | 45.449 | 3442.061 | < 0.01 |
| Developmental temperature | 1 | 0.006 | 0.006 | 0.456 | 0.503 |
| Immediate temperature | 1 | 1.715 | 1.715 | 129.910 | < 0.01 |
| Developmental temperature*Immediate temperature | 1 | 0.130 | 0.130 | 9.844 | < 0.01 |
| Error | 52 | 0.687 | 0.013 |  |  |
| **Number of chirps / 10 s** |  |  |  |  |  |
| Intercept | 1 | 7020.214 | 7020.214 | 3104.616 | < 0.01 |
| Developmental temperature | 1 | 1.929 | 1.929 | 0.853 | 0.360 |
| Immediate temperature | 1 | 242.881 | 242.881 | 107.412 | < 0.01 |
| Developmental temperature*Immediate temperature | 1 | 22.881 | 22.881 | 10.119 | < 0.01 |
| Error | 52 | 117.583 | 2.261 |  |  |

| **Syllable duration (s)** | **Degree of Freedom** | **SS** | **MS** | **F** | **P** |
| --- | --- | --- | --- | --- | --- |
| Intercept | 1 | 0.012 | 0.012 | 5630.228 | < **0.01** |
| Developmental temperature | 1 | 0.000 | 0.000 | 5.389 | **0.024** |
| Immediate temperature | 1 | 0.000 | 0.000 | 7.744 | < **0.01** |
| Developmental temperature*Immediate temperature | 1 | 0.000 | 0.000 | 5.302 | **0.025** |
| Error | 52 | 0.000 | 0.000 |  |  |
| **Syllable period (s)** | **Degree of Freedom** | **SS** | **MS** | **F** | **P** |
| Intercept | 1 | 0.038 | 0.038 | 9063.547 | < **0.01** |
| Developmental temperature | 1 | 0.000 | 0.000 | 4.483 | **0.039** |
| Immediate temperature | 1 | 0.000 | 0.000 | 61.881 | < **0.01** |
| Developmental temperature *Immediate temperature | 1 | 0.000 | 0.000 | 12.906 | < **0.01** |
| Error | 52 | 0.000 | 0.000 |  |  |
| **Number of syllables/chirp** | **Degree of Freedom** | **SS** | **MS** | **F** | **P** |
| Intercept | 1 | 11590.960 | 11590.960 | 3578.903 | < **0.01** |
| Developmental temperature | 1 | 1.960 | 1.960 | 0.605 | 0.440 |
| Immediate temperature | 1 | 0.350 | 0.350 | 0.108 | 0.744 |
| Developmental temperature *Immediate temperature | 1 | 5.630 | 5.630 | 1.738 | 0.193 |
| Error | 52 | 168.410 | 3.240 |  |  |
| **Peak frequency (Hz)** | **Degree of Freedom** | **SS** | **MS** | **F** | **P** |
| Intercept | 1 | 1265604000 | 1265604000 | 42380.200 | < **0.01** |
| Developmental temperature | 1 | 568138 | 568138 | 19.020 | < **0.01** |
| Immediate temperature | 1 | 154106 | 154106 | 5.160 | **0.027** |
| Developmental temperature*Immediate temperature | 1 | 32252 | 32252 | 1.080 | 0.304 |
| Error | 52 | 1552882 | 29863 |  |  |

## 4. Discussion

Our study provides conclusive evidence of the profound impacts of temperature on the life-history, body morphometry and various features of the mating call of *A. asiaticus*. Individuals raised at the higher temperature reached adulthood faster but had reduced lifespan than individuals raised at the lower temperature. *Acanthogryllus asiaticus* was also found to be an exception to the temperature-size rule as body size parameters were found to be higher for males raised at the higher temperature. In addition, both immediate ambient temperature and developmental temperature independently and interactively influenced the mating call of *A. asiaticus*.

### 4.1. Effect of temperature on life-history traits

Our results show that individuals raised at 30°C showed rapid development and reached adulthood 75 days faster than the individuals at 25°C. We also found that the adult lifespan of individuals raised at 25°C was higher than those at 30°C. This result suggests that higher temperature supports faster development but not a longer lifespan. A similar trend has been observed in *Teleogryllus emma*, where individuals raised at 35°C reached adulthood 80 days faster but lived shorter than the individuals at lower temperatures (Kim et al., 2007). Likewise, in *G. bimaculatus*, *Acheta domesticus* and *G. texensis*, the developmental time decreased with increasing temperature (Behrens et al., 1983; Booth and Kiddell, 2007; Adamo and Lovett, 2011). Studies on other insect models have also reported a similar trend. For example, in 4 species of *Culex* mosquitoes, an increase of temperature from 16-24°C resulted in an average 2.9-fold increase in developmental rate (Ciota et al., 2014).

This trend of rapid development at increased temperature can be explained by the metabolic theory of ecology (Gillooly et al., 2001) which posits that metabolic rate increases almost exponentially with temperature and since developmental time is dependent on metabolic rate, it decreases with temperature. Despite the increased metabolic rate in higher temperature, the increased developmental time in lower temperature can result in an increased energy cost of development as reported in the lizard *Sceloporus undulates* (Angilletta et al., 2000). In *A. domesticus*, Booth and Kiddell (2007) found energy expenditure to be higher at 25°C compared to 28°C and suggested that 25°C is a sub-optimal temperature. In *A. asiaticus*, energy expenditure at 25 and 30°C remains to be investigated, however, slower development at 25°C suggests that 25°C is likely the sub-optimal temperature for *A. asiaticus*.

According to temperature-size rule (Atkinson, 1994), it was expected that the size of *A. asiaticus* would decrease with an increase in temperature. We found mixed results when we examined the relationship between temperature and body size. In our study, pronotum length and width of males and ovipositor length of females follow the converse of temperature-size rule (Atkinson, 1994), whereas the body length and wing size of females abide by this rule. While studies examining the effect of temperature on body size are very limited, this inconsistency with the temperature-size rule has also been confirmed in other cricket species. For instance, Roe et al. (1985) showed that individuals of *A. domesticus* raised at 35°C had a higher dry mass than those raised at 25°C. However, on the same species, Booth and Kiddell (2007) found that individuals at 25°C had a higher dry mass than those at 28°C. Such contrasting results could be because of the differences in adult dry mass for the species reported by the two studies. Another study on *G. firmus* showed that the trend was not consistent across all the examined parameters except for ovipositor, which did not follow the temperature-size rule (Bégin et al., 2004). Studies on other ectotherms have also reported exceptions to the temperature-size rule. For instance, adult mass in temperate grasshopper, *Chorthippus brunneus*, also did not follow the temperature-size rule (Walters and Hassall, 2006). Walters and Hassall (2006) suggested that the lower temperature threshold for development than growth could be one of the mechanistic reasons for the deviation from the temperature-size rule. This opens avenues for detailed physiological studies in *A. asiaticus* to test these predictions.

### 4.2. Effect of temperature on mating call features

Our study suggests that immediate temperatures (22-30°C) influence different call features of field-collected males from the same season. The signals recorded at 30°C had the maximum chirp rate and peak frequency, but all other call features were minimum at this temperature. The relationship between chirp rate and temperature has been described as Dolbear’s law (Dolbear, 1987). For instance, in *T. oceanicus* and *G. bimaculatus*, the chirp rate increased linearly with temperature (Walker and Cade, 2003; Doherty, 1985). Similar to the relationship between temperature and chirp rate, the display rate of various signals of other ectotherms is also influenced by temperature. For instance, the pulse rate of *Bufo americanus* toad call increased linearly with temperature (Zweifel, 1968) whereas, the rate of stridulatory scrapes of *Habronattus clypeatus* spider increased to a point and then decreased (Brandt et al., 2018). In our species, the frequency increased with temperature, but the effect of temperature on peak frequency does not follow a general trend across cricket species. For instance, the peak frequency increased by 400 Hz in *G. integer* and 1500 Hz in *Plebeiogryllus guttiventris* in response to an increase of 12°C and 16°C respectively (Martin et al., 2000; Mhatre and Balakrishnan, 2006). However, studies on *G. firmus* and *G. bimaculatus* showed that an increase in temperature does not affect the peak frequency (Pires and Hoy 1992; Doherty 1985). The absence of a general trend might be because of the individual differences among males in the rate of closing wing stroke and wing mass which influence peak frequency and mask the effect of temperature (Walker 1962; Martin et al., 2000). The effect of temperature on different call features can be attributed to the constraints it poses on physiological and biochemical factors involved in muscle function, which influences motor activities responsible for sound production (Greenfield and Medlock, 2007). Since the immediate temperature affects signalling, animals can exhibit preferences for the temperature of the display site, which may drive microhabitat selection. For instance, *G. integer* males showed a preference for warmer open cracks (Hedrick et al., 2002) and males of *Hyla versicolor* called from a warmer environment in all seasons (Höbel and Barta, 2014). Preference for the temperature of the calling site is yet to be investigated in our species.

Our results also demonstrate the impact of developmental temperature on the mating signals of crickets, thereby highlighting the importance of rearing microclimate on the fitness of the organism. Individuals raised at 30°C called at a higher peak frequency than the individuals raised at 25°C. Moreover, we found that the interaction between developmental temperature and immediate ambient temperature influences different call features differentially. The effect of developmental temperature has been examined on the other cricket species. For instance, the study on *L. cerasina* showed that the males reared at 25°C called at a faster pulse rate than those reared at 20°C, but the peak frequency did not differ between the two groups (Grace and Show, 2004). In *G. rubens,* the individuals from the fall season called at a faster rate and higher peak frequency than those from the spring season (Beckers et al., 2019). However, in both these studies, the call recordings were done at only one temperature; 20 and 24°C, respectively (Grace and Shaw, 2004; Beckers et al., 2019). Similar to our study, Olvido and Mousseau (1995) tested the effect of two different developmental environments (31°C, 15L:9D and 24°C, 11L:13D) in striped ground cricket, *Allonemobius fasciatus*, by recording the individuals at three different ambient temperatures (24, 28 and 31°C). They found that chirp rate, chirp duration, inter-chirp interval, pulse number, and carrier frequency were affected by both developmental and immediate temperature. Therefore, both our study and study by Olvido and Mousseau (1995) suggest that the developmental effect on call parameters vary with the immediate calling environment as different developmental environments can lead to inconsistent changes in call features in different immediate calling environments.

The influence of developmental and immediate ambient temperature on call features might alter the attractiveness of the call. We found that with higher temperature, chirp rate increased and the chirp period decreased. Given that female crickets are known to prefer calls with higher chirp rates and longer chirps (Wagner, 1996), it is expected that calls produced by individuals at a higher temperature are more favoured than those at a lower temperature. However, the impact of temperature on signalling can also affect the efficacy of sexual communication, which can be resolved by a phenomenon known as temperature coupling (Gerhardt, 1978). This phenomenon suggests a parallel shift in male signals and female preferences in a short◻term and reversible fashion in response to the immediate ambient temperature and has been reported in a group of insects (Doherty and Hoy, 1985; Pires and Hoy,1992) and anurans (Gerhardt, 1978). So far, such parallel plasticity of signals and signal preferences for the developmental temperature have been reported in the Hawaiian cricket, *L. cerasina* (Grace and Shaw, 2004) and *G. rubens* (Beckers et al., 2019). Our findings with *A. asiaticus* indicate that immediate and developmental temperature both independently and interactively can lead to differences in call features. This plasticity in calls opens up the potential for examining temperature coupling and the implications of this in the mate choice of this bivoltine species.

In conclusion, our study reveals that developmental temperature appears to impact the life-history of a nocturnal ectotherm which may have major consequences on their fitness. It also provides insights regarding the ecological and evolutionary consequences of temperature rise on intersexual communication. In addition, understanding the influence of developmental environment on various life functions of tropical insects can be crucial for predicting their responses to climate change.

## Acknowledgements

The research was supported by grants from IISER Mohali to MJ; RS was supported by INSPIRE fellowship and PP was supported by INSPIRE SHE scholarship by Department of Science & Technology, India, (https://online-inspire.gov.in/). We thank Gurmeet Singh for help with lab-culture maintenance.

## Author contributions

MJ conceived the study; MJ and RS designed the experiments; RS performed the experiments, collected raw data and conducted signal analyses with PP’s help. RS and MJ carried out statistical analyses and wrote the manuscript; PP helped with editing and finalizing it. All authors approved the final version of the manuscript and are accountable for the content therein.

