## Supplementary File for "Effect of temperature on life-history traits and mating calls of a field cricket, *Acanthogryllus asiaticus*"

Manjari Jain

Department of Biological Sciences,

Indian Institute of Science Education and Research, Mohali,

Punjab 140306, India

**Abstract**

Ectotherms are sensitive to the changes in ambient temperature with respect to their physiology and development. To compensate for the effects of variation in temperature, ectotherms exhibit physiological plasticity which can be for short or long term. An extensive body of literature exists towards understanding these effects and the solutions ectotherms have evolved. However, to what extent rearing temperature during early life stages impacts the behaviour expressed in adulthood is less clearly understood. In the present study, we aimed to examine the effect of developmental temperature on life-history traits and mating call features in a tropical field cricket, *Acanthogryllus asiaticus*. We raised *A. asiaticus* at two different developmental conditions: 25°C and 30°C. We found developmental time and adult lifespan of individuals reared at 30°C to be shorter than those at 25°C. Increased developmental temperature influenced various body size parameters differentially. Males raised at 30°C were found to be larger and heavier than those raised at 25°C, making *A. asiaticus* an exception to the temperature-size rule. We found a significant effect of the change in immediate ambient temperature on different call features of both field-caught and lab-bred individuals. In addition, developmental temperature also affected mating call features as individuals raised at higher temperature produced faster calls with a higher peak frequency compared to those raised at lower temperature. However, the interaction of both developmental and immediate temperature on mating calls showed differential effects. Our study highlights the importance of understanding how environmental temperature shapes life-history and sexual communication in crickets.

Appendix S1

Rearing: Segregated eggs were placed on wet cotton pads in a petri-dish which was kept in plastic containers (15 X 12 X 10 cm) having lids with a 10 X 5 cm opening covered with mosquito screening mesh to allow air circulation. Once hatched, nymphs were transferred to larger plastic containers (35 X 25 X 12 cm) having lids with mesh-covered opening. The bottom of each of this container was lined with egg cartons and two petri-dishes filled with dogfood powder and wet cotton pads. Periodic observations were carried out every day till the time of nymph appearance and then after every three days. After the final moult, each individual was kept in a separate box of diameter 12 cm and height 5 cm with a mesh (9 X 8 cm) lid and food and water was provided ad libitum.

**Table S1** Comparison of different life-history traits of *Acanthogryllus asiaticus* at 25°C and 30°C using t test. Significant values are indicated in bold.

| | Mean ± SD | Mean ± SD | t-value | df | P | N | N |
|---|---|---|---|---|---|---|---|
| (days) | 25℃ | 30℃ | | | | 25℃ | 30℃ |
| Nymph appearance | 22.37 ± 5.26 | 10.89 ±3.87 | 20.57 | 271 | **<0.01** | 135 | 138 |
| Developmental time | 170.5± 24.67 | 95.87 ± 27.85 | 10.29 | 52 | **<0.01** | 24 | 30 |
| Adult lifespan | 74.5 ± 18.36 | 59.07 ± 24.81 | 2.54 | 52 | **<0.01** | 24 | 30 |

**Table S2** Comparison of body size parameters in males and females of *Acanthogryllus asiaticus* at 25°C and 30°C using t test. Significant values are indicated in bold.

| | Mean ± SD | Mean ± SD | t-value | df | P |
|---|---|---|---|---|---|
| **Female** | 25℃ | 30℃ | | | |
| Weight (g) | 0.28 ± 0.04 | 0.26 ± 0.05 | 0.709 | 13 | 0.491 |
| Body length (mm) | 17.82 ± 1.46 | 14.82 ± 1.03 | 5.116 | 16 | **<0.01** |
| Pronotum length (mm) | 2.60 ± 0.15 | 2.74 ± 0.20 | -1.669 | 16 | 0.114 |
| Pronotum width (mm) | 4.45 ± 0.17 | 4.57 ± 0.16 | -1.485 | 16 | 0.157 |
| Wing length (mm) | 9.58 ± 0.49 | 8.85 ± 0.48 | 3.157 | 16 | **<0.01** |
| Ovipositor length (mm) | 5.31 ± 0.28 | 5.76 ± 0.20 | 3.981 | 16 | **<0.01** |

| | | | | | |
|---|---|---|---|---|---|
| **Male** | | | | | |
| Weight (g) | 0.21 ± 0.04 | 0.24±0.03 | -2.292 | 32 | **0.027** |
| Body length (mm) | 15.43 ± 1.23 | 15.72 ± 0.96 | -0.842 | 38 | 0.405 |
| Pronotum length (mm) | 2.45 ± 0.26 | 2.69 ± 0.17 | -3.488 | 38 | **<0.01** |
| Pronotum width (mm) | 4.38 ± 0.31 | 4.69 ± 0.25 | -3.384 | 38 | **<0.01** |
| Wing length (mm) | 9.06 ± 0.58 | 9.21 ± 0.58 | -0.825 | 38 | 0.415 |

**Table S3 A.** Comparison of call properties when of individuals reared at 25°C and recorded at different temperatures (22, 24, 26, 28 and 30°C) during short term exposure using Kruskal-Wallis Anova. **B.** Descriptive statistics for all the call features recorded at different temperature. **C.** Pairwise comparison of call parameters recorded at different temperatures using Mann-Whitney test with Bonferroni correction. Significant differences indicated in bold.

| **A. Call properties** | **H** | **df** | **N** | **P** |
|---|---|---|---|---|
| Chirp duration (s) | 152.856 | 4 | 362 | **<0.01** |
| Chirp period (s) | 111.367 | 4 | 347 | **<0.01** |
| Syllable duration (s) | 394.718 | 4 | 4644 | **<0.01** |
| Syllable period (s) | 607.075 | 4 | 4283 | **<0.01** |
| Peak frequency (Hz) | 76.346 | 4 | 422 | **<0.01** |
| No of syllables per chirps | 52.129 | 4 | 362 | **<0.01** |
| No of chirps per 10s | 26.900 | 4 | 71 | **<0.01** |

| **B. Mean ± SD** | **22°C** | **24°C** | **26°C** | **28°C** | **30°C** |
|---|---|---|---|---|---|
| Chirp duration (s) | 0.34 ± 0.06 | 0.35 ± 0.04 | 0.38 ± 0.03 | 0.31 ± 0.03 | 0.28 ± 0.04 |
| Chirp period (s) | 0.95 ± 0.17 | 0.94 ± 0.12 | 0.92 ± 0.12 | 0.90 ± 0.13 | 0.73 ± 0.12 |
| Peak frequency (Hz) | 4234.04 ± 126 | 4059.71 ± 262 | 4327.22 ± 163 | 4370.62 ± 221 | 4384.16 ± 216 |
| Syllable duration (s) | 0.015 ± 0.003 | 0.014 ± 0.003 | 0.017 ± 0.005 | 0.015 ± 0.005 | 0.013 ± 0.003 |
| Syllable period (s) | 0.027 ± 0.007 | 0.027 ± 0.007 | 0.028 ± 0.007 | 0.025 ± 0.007 | 0.022 ± 0.007 |
| No of syllables per chirps | 12.59 ± 2.14 | 12.93 ± 1.51 | 14.44 ± 1.11 | 13.29 ± 1.33 | 12.57 ± 1.47 |
| No of chirps per 10s | 10.86 ±1.51 | 11.54 ± 1.20 | 11.27 ± 1.42 | 11.33 ± 0.82 | 14.00 ± 1.68 |

| **C.** Comparison | Chirp duration (s) | Chirp period (s) | Syllable duration (s) | Syllable period (s) | Peak frequency (Hz) | No of syllables per chirps | No of chirps per 10 s |
|---|---|---|---|---|---|---|---|
| 22°C vs 24°C | ns | ns | **<0.01** | **<0.01** | **<0.01** | ns | ns |
| 22°C vs 26°C | **<0.01** | ns | **<0.01** | ns | **<0.01** | **<0.01** | ns |
| 22°C vs 28°C | **<0.01** | ns | **<0.01** | **<0.01** | **<0.01** | **<0.01** | ns |
| 22°C vs 30°C | **<0.01** | **<0.01** | **<0.01** | **<0.01** | **<0.01** | ns | **<0.01** |
| 24°C vs 26°C | **<0.01** | ns | **<0.01** | **<0.01** | **<0.01** | **<0.01** | ns |
| 24°C vs 28°C | **<0.01** | ns | **<0.01** | **<0.01** | **<0.01** | ns | ns |
| 24°C vs 30°C | **<0.01** | **<0.01** | **<0.01** | **<0.01** | **<0.01** | ns | **<0.01** |
| 26°C vs 28°C | **<0.01** | ns | **<0.01** | **<0.01** | ns | **<0.01** | ns |
| 26°C vs 30°C | **<0.01** | **<0.01** | **<0.01** | **<0.01** | ns | **<0.01** | **<0.01** |
| 28°C vs 30°C | **<0.01** | **<0.01** | **<0.01** | **<0.01** | ns | **<0.01** | **<0.01** |

**Table S4.** Comparison of call features of individuals bred at 25℃ and recorded at 25℃ and 30℃ using t – test to examine the effect of immediate temperature. Significant differences indicated in bold.

| Call parameters | Mean ± SD 25°C | Mean ± SD 30°C | t-value | df | P | N 25°C | N 30°C |
|---|---|---|---|---|---|---|---|
| Chirp duration (s) | 0.42 ± 0.07 | 0.36 ± 0.05 | 2.34 | 22 | **0.03** | 12 | 12 |
| Chirp period (s) | 1.03 ± 0.12 | 0.77 ± 0.07 | 6.22 | 22 | **<0.01** | 12 | 12 |
| Number of chirps/10 s | 9.67 ± 1.15 | 12.58 ± 1.38 | -5.62 | 22 | **<0.01** | 12 | 12 |
| Syllable duration (s) | 0.015 ± 0.001 | 0.015 ± 0.001 | 0.32 | 22 | 0.75 | 12 | 12 |
| Syllable period (s) | 0.028 ±0.002 | 0.026 ± 0.002 | 2.42 | 22 | **0.02** | 12 | 12 |
| Number of syllables/chirp | 15.13 ± 2.03 | 14.33 ± 1.23 | 1.17 | 22 | 0.25 | 12 | 12 |
| Peak frequency (Hz) | 4672.70 ± 190.56 | 4730.21 ± 207.41 | -0.71 | 22 | 0.48 | 12 | 12 |

**Table S5.** Comparison of call features of individuals bred at 30℃ and recorded at 25℃ and 30℃ using t – test to examine the effect of immediate temperature. Significant differences indicated in bold.

| | Mean ± SD 25°C | Mean ± SD 30°C | t-value | df | P | N 25°C | N 30°C |
|---|---|---|---|---|---|---|---|
| Chirp duration (s) | 0.40 ± 0.06 | 0.33 ± 0.05 | 3.62 | 30 | **<0.01** | 16 | 16 |
| Chirp period (s) | 1.15 ± 0.15 | 0.70 ±0.08 | 10.28 | 30 | **<0.01** | 16 | 16 |
| Number of chirps/10 s | 8.75 ± 1.48 | 14.25±1.81 | -9.41 | 30 | **<0.01** | 16 | 16 |
| Syllable duration (s) | 0.015 ± 0.001 | 0.013 ± 0.001 | 3.84 | 30 | **<0.01** | 16 | 16 |
| Syllable period (s) | 0.029 ± 0.002 | 0.022 ± 0.001 | 10.24 | 30 | **<0.01** | 16 | 16 |
| Number of syllables/chirp | 14.11 ± 1.95 | 14.59 ± 1.8 | -0.72 | 30 | 0.48 | 16 | 16 |
| Peak frequency (Hz) | 4827.74 ± 148.2 | 4982.24 ± 152.9 | -2.90 | 30 | **<0.01** | 16 | 16 |

**Table S6.** Comparison of call features of individuals bred at 25°C and 30°C and recorded at 30°C using t – test to examine the effect of developmental temperature. Significant differences indicated in bold.

| **Call parameters** | **Mean ± SD 25°C** | **Mean ± SD 30°C** | **t-value** | **df** | **P** | **N 25°C** | **N 30°C** |
|---|---|---|---|---|---|---|---|
| Chirp duration (s) | 0.36 ± 0.05 | 0.33 ± 0.05 | 1.855 | 26 | 0.08 | 12 | 16 |
| Chirp period (s) | 0.77 ± 0.07 | 0.70 ± 0.08 | 2.468 | 26 | **0.02** | 12 | 16 |
| Number of chirps/10 s | 12.58 ±1.4 | 14.25 ± 1.81 | -2.661 | 26 | **0.01** | 12 | 16 |
| Syllable duration (s) | 0.015 ± 0.001 | 0.013 ± 0.001 | 3.769 | 26 | **<0.01** | 12 | 16 |
| Syllable period (s) | 0.026 ± 0.003 | 0.022 ± 0.001 | 4.168 | 26 | **<0.01** | 12 | 16 |
| Number of syllables/chirp | 14.33 ± 1.23 | 14.59 ± 1.81 | -0.432 | 26 | 0.67 | 12 | 16 |
| Peak frequency (Hz) | 4730.206 ± 207.4 | 4982.236 ±152.9 | -3.707 | 26 | **<0.01** | 12 | 16 |

**Table S7.** Comparison of call features of individuals bred at 25°C and 30°C and recorded at 25°C using t – test to examine the effect of developmental temperature. Significant differences indicated in bold.

| **Call parameters** | **Mean ± SD 25°C** | **Mean ± SD 30°C** | **t-value** | **df** | **P** | **N 25°C** | **N 30°C** |
|---|---|---|---|---|---|---|---|
| Chirp duration (s) | 0.42 ± 0.07 | 0.40 ± 0.06 | 0.971 | 26 | 0.340 | 12 | 16 |
| Chirp period (s) | 1.03 ± 0.12 | 1.15 ± 0.15 | -2.200 | 26 | **0.037** | 12 | 16 |
| Number of chirps/10 s | 9.67 ± 1.15 | 8.75 ± 1.48 | 1.773 | 26 | 0.088 | 12 | 16 |
| Syllable duration (s) | 0.015 ± 0.001 | 0.015 ± 0.001 | 0.012 | 26 | 0.991 | 12 | 16 |
| Syllable period (s) | 0.028 ±0.002 | 0.029 ± 0.002 | -1.012 | 26 | 0.321 | 12 | 16 |
| Number of syllables/chirp | 15.13 ± 2.03 | 14.11 ± 1.95 | 1.343 | 26 | 0.191 | 12 | 16 |
| Peak frequency (Hz) | 4672.70 ± 190.56 | 4827.738 ± 148.2 | -2.425 | 26 | **0.023** | 12 | 16 |
